# Targeted protein depletion in quiescent fission yeast cells using an auxin-degron system

**DOI:** 10.64898/2026.11.20.767346

**Authors:** Denez Le Collen, Alicia Vázquez-Bolado, Pei-Yun Jenny Wu

## Abstract

Quiescence is a reversible, non-proliferative cellular state in which cells can remain for extended periods of time while retaining the ability to re-enter the cell cycle upon exposure to favorable conditions. This physiological state is critical for a broad range of biological processes, from stress response in unicellular systems to differentiation and development in multicellular organisms. Surprisingly, despite its recognized roles in cell biology in diverse species, key aspects of the regulation of quiescence remain poorly understood. To investigate protein function specifically in quiescence, a conditional activation approach is critical, in particular for factors that are essential for cell viability. However, the efficacy of a number of such inactivation strategies in quiescent cells remained unknown. Using nitrogen-depleted fission yeast cells as a well-established model for quiescence, we probed two types of commonly-used approaches: a targeted method for the reversible inhibition of engineered kinases using non-hydrolyzable ATP analogs, and a more general system targeting protein depletion using an auxin-inducible degron. Here we show that the use of analog-sensitive kinases is unsuitable for quiescent fission yeast cells, as the ATP analog serves as a nitrogen source and induces cell cycle re-entry. We then demonstrate how coupling a degron tag with a specific auxin ligand allows for rapid, lasting and reversible protein depletion in quiescent cells, providing a powerful tool for investigating the regulation of this crucial and widespread cellular state.

**Take Away:**

- A synthetic auxin/degron system induces fast and long-lasting protein depletion in quiescent fission yeast cells.
- Synthetic auxin-induced degradation in quiescence can be rapidly reversed upon ligand removal.
- The analog-kinase inhibition approach is inadvisable for nitrogen-starved cells.

## Introduction

Quiescence is a prevalent cellular state in nature, and most cells spend a large part of their life cycle in non-dividing states. Remarkably, quiescent cells remain metabolically active for extended periods of time, maintaining their ability to resume growth and proliferation in response to appropriate signals or environmental cues. In multicellular organisms, quiescence is crucial for a variety of processes ranging from development to immunity (de Morree & Rando, 2023). In unicellular species, quiescence allows cells to resist environmental stresses such as nutrient deprivation and temperature fluctuations (Su et al., 1996; Sun & Gresham, 2021). Surprisingly, although this cellular state is integral to the biology of living organisms, much remains unknown about the molecular mechanisms governing quiescence.

The fission yeast *Schizosaccharomyces pombe* is a unicellular eukaryote that is widely used to investigate fundamental cell biological processes. In particular, it is amenable to advanced genetic engineering and shares a number of essential features with more complex eukaryotes, from cell cycle control to the regulation of chromatin and gene expression (Vyas et al., 2021). In fission yeast, quiescence can be induced by nutrient deprivation, including glucose or nitrogen starvation (Yanagida, 2009). Compared to proliferative cells, nitrogen-starved cells are smaller (Fig. 1A), show alterations in chromatin organization, harbor lower RNA and protein levels, and maintain their capacity to re-renter the cell cycle for long periods of time (Cai et al., 2020; Marguerat et al., 2012; Su et al., 1996). These characteristics are consistent with what has been observed in quiescent cells in more complex eukaryotes (Braun et al., 2026; de Morree & Rando, 2023), making nitrogen-depleted fission yeast cells a model of choice for investigating the core mechanisms regulating this cellular state.

**Figure 1.**
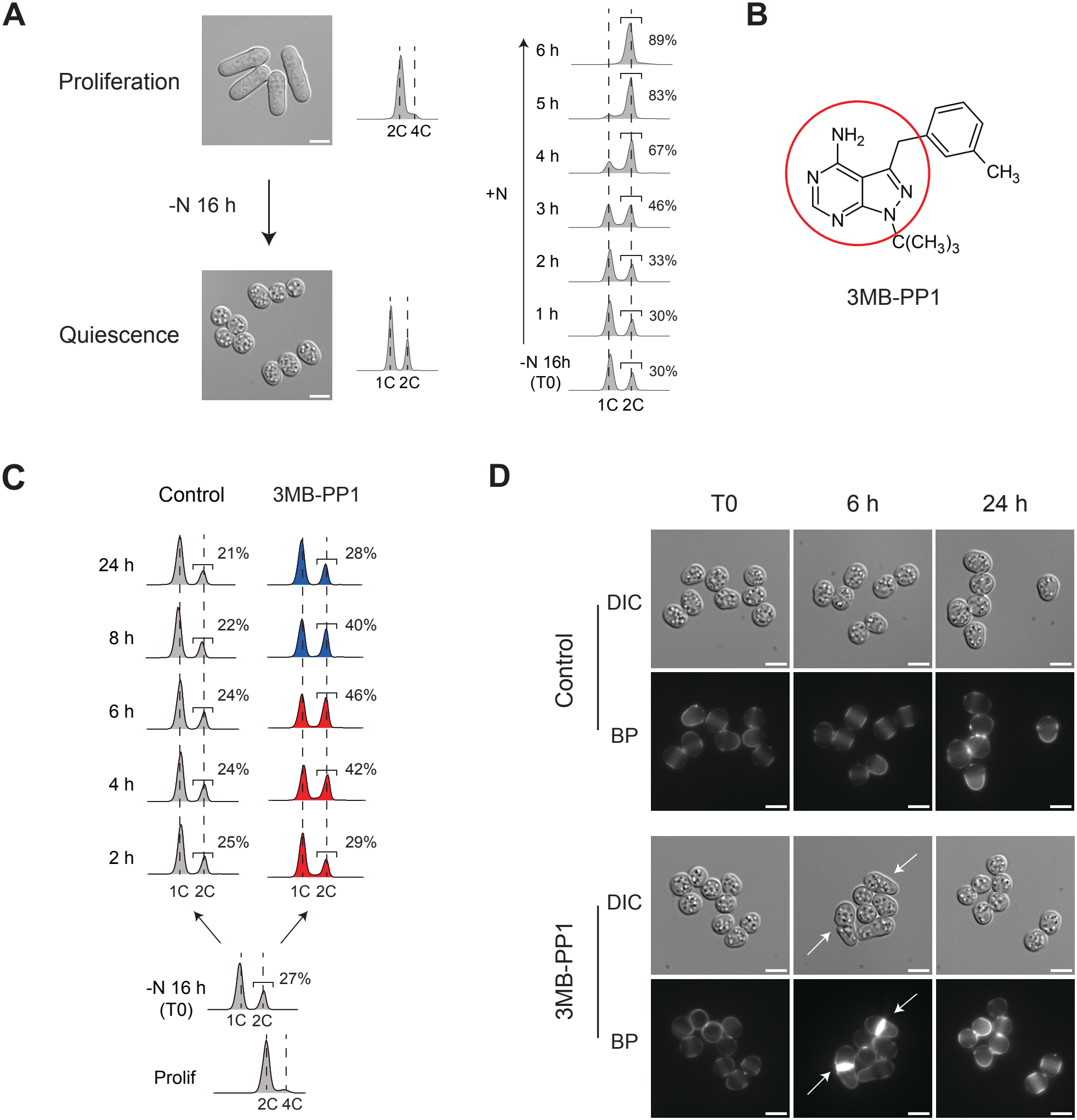
The 3MB-PP1 ATP analogue triggers cell cycle re-entry in nitrogen-starved quiescent cells. **A**. In fission yeast cells, nitrogen depletion induces reversible cell cycle exit and entry into quiescence. Left: Proliferating cells shifted to nitrogen-depleted medium for 16 h exit the division cycle and enter quiescence at small sizes and with a 1C DNA content. Upon addition of nitrogen, these cells (-N 16 h/T0) undergo DNA replication followed by cell division. DIC images of both proliferating and quiescent cells are shown. Scale bar = 5µm. Right: Flow cytometry analysis of DNA content during cell cycle re-entry. Percentages of cells with 2C DNA content are indicated. **B**. Chemical structure of 3MB-PP1, a non-hydrolyzable ATP analog that is commonly used for the dose-dependent and reversible inhibition of engineered kinases (C. Zhang et al., 2013). Red circle highlights the nitrogen groups. **C**. DNA content analysis of wild-type quiescent cells treated with DMSO (Control) or 20 μM 3MB-PP1 for the indicated times. Red profiles: a subset of cells undergo DNA replication, as shown by an increase in the percentages of 2C cells. Blue profile: after DNA replication, these cells undergo division (see also *D*) and re-enter quiescence with a 1C DNA content. Prolif: proliferating cells prior to nitrogen starvation. -N 16 h/T0: nitrogen-starved quiescent cells. **D**. Representative images of cells in *C* at the indicated time points. DIC: Differential interference contrast. BP: Blankophor. Arrows indicate dividing cells. Scale bars = 5µm.

In this context, gene deletion studies have provided valuable insight into key factors involved in quiescence maintenance (Roche et al., 2016; Sajiki et al., 2018; Sideri et al., 2014; Zahedi et al., 2020). However, assessing the role of proteins in the control of quiescence, in particular when they are essential for proliferation, requires conditional approaches that can be triggered specifically in quiescent cells, after cell cycle exit has taken place. Although temperature-sensitive mutants have contributed to the identification of a number of regulators of quiescence (Sajiki et al., 2009), this strategy combines temperature and nutritional stresses, making it more difficult to ascertain precisely the roles of candidate genes in quiescence. In addition, as most alternative conditional techniques for the downregulation of protein function have been developed for proliferating cells, their effectiveness in this non-dividing state remains to be determined.

In this study, we have tested two different strategies for inducing functional inhibition of proteins in nitrogen-depleted quiescent cells. First, we focused on the targeted alteration of kinase activities, which are central to a number of processes in both proliferative and quiescent states. Second, we evaluated a more general approach, in which protein degradation is specifically induced in quiescent cells by an auxin-degron system. Interestingly, we showed that the use of engineered analog-sensitive kinases is unsuitable for nitrogen-starved quiescent cells, as the non-hydrolyzable ATP analog serves as a nitrogen source. Importantly, we then demonstrated the power of the auxin-induced degradation method when coupled with a specific auxin ligand, which enabled rapid and reversible protein depletion in quiescent cells. Our study therefore provides an efficient tool that can be applied for functional downregulation in non-proliferating cells, opening the door to uncovering key regulators of quiescence.

## Materials and Methods

### Fission yeast strains and methods

Fission yeast strains used in this study are listed in Table 1. Standard *S. pombe* genetic procedures and media were used (Moreno et al., 1991). Cells were grown in minimal medium plus supplements (EMM6S) unless otherwise indicated. YE4S: nutrient-rich medium. To induce quiescence by nitrogen depletion, cultures were grown at 25 °C to a density of ∼2 × 10^6^ cells in EMM6S. Cells were then washed 3 times and resuspended in minimal medium without NH_4_Cl (EMM-N) for 16 h at 25 °C, leading to cell cycle exit. Release from starvation was induced by addition of 0.5 g/L NH_4_Cl.

**Table 1.** Strains used in this study.

| **Number** | **Genotype** |  |
|---|---|---|
| JW1575 | *h+ rpb1-HA-ura* | *This study* |
| JW2488 | *h+ rpb1-HA-ura seb1::3xsAID::KanMx6 urg3:OsTIR:urg1* | *This study* |
| JW2491 | *h- rpb1-HA-ura seb1::3xsAID::KanMx6* | *This study* |
| JW2497 | *h+ urg3::OsTIR1::urg1* | *This study* |
| JW2535 | *h+ rpb1-HA-ura dis2::3xsAID::KanMx6 urg3:OsTIR:urg1* | *This study* |
| JW2536 | *h- rpb1-HA-ura dis2::3xsAID::KanMx6* | *This study* |

A prototrophic strain harboring *rpb1-HA* (JW1575) was constructed by crossing FY17150 (*JY741 h-rpb1-HA ura4+ ade6-M216 leu1*; National BioResource Project (NBRP Yeast), Japan, JPNBRP202225) with PN1 (*h-972*). For *seb1* and *dis2* C-terminal tagging, integration sites were directed by primers containing 80-bp homology to regions flanking the stop codon, and 3xsAID was PCR-amplified from *pFA6a-3×sAID-kanMX* (X.-R. Zhang et al., 2022) . To generate strains containing *OsTIR1* at the *urg3-urg1* locus, CRISPR editing was performed following the SpEDIT protocol (Torres-Garcia et al., 2020). Briefly, a 20 bp sgRNA sequence was cloned into the pLSB-NAT plasmid using Golden Gate assembly. For *OsTIR* integration, the repair fragment was amplified by PCR from a plasmid containing *Padh1-OsTIR-tadh1* (X.-R. Zhang et al., 2022) using oligonucleotides with homology to regions covering the insertion site, located between *urg*3 and *urg1*. Cells were then co-transformed with the sgRNA/Cas9 plasmid and the repair template.

For treatment with 3MB-PP1, a 10 mM stock solution of 3MB-PP1 (4-amino-1-tert-butyl-3-(3-methylbenzyl)pyrazolo[3,4-d]pyrimidine, TRC-A602960, LGC Group) was prepared in DMSO. Cells were treated with 20 µM 3MB-PP1, with control cultures being exposed to an equivalent volume of DMSO.

For treatment with NAA, 0.5 M stock solution of NAA (1-Naphthaleneacetic acid; Apollo Scientific) was prepared in 1 M NaOH. Cells were treated with 0.5 mM NAA, with control cultures being exposed to an equivalent volume of 1M NaOH.

### DNA content analysis by flow cytometry

Cells were fixed with 1 mL of 70% cold ethanol and stored at 4°C. Cells were then washed with 50 mM sodium citrate, followed by overnight incubation at 37 ºC with RNaseA (0.1 mg/mL, Sigma-Aldrich). DNA was stained with 1 µg/mL propidium iodide (Sigma-Aldrich), and cells were briefly sonicated using a Branson Digital Sonifier. Samples were processed using a BD Accuri C6 (BD Biosciences) flow cytometer. Data were analyzed using FlowJo (BD Biosciences). For Figures 1A, 1C and 2C, the percentage of cells with 2C DNA content was determined by taking the total number of 1C and 2C cells in a population, then calculating the proportion of 2C cells.

### Microscopy

Images were acquired using a Zeiss Axio Observer epifluorescence microscope (Zeiss GmbH) equipped with a Lumencor Spectra X illumination system (Lumencor), an Orca Flash 4.0V2 sCMOS camera (Hamamatsu Photonics) and the VisiView acquisition software (Visitron Systems GmbH). For visualizing division septa, live cells were stained with 0.5 mg/mL Blankophor (Ambeed Inc.).

### Spot assays

For Figures 2E and 3D, cells were depleted of nitrogen for 16 h to enter quiescence and spotted at 0, 1, and 4 days after NAA or NaOH treatment. Cells were serially diluted tenfold, spotted onto YE4S plates and incubated at 25 °C. For Figure 3B, proliferating cells of the indicated genotypes were spotted on YE4S plates containing 0.5 mM NAA (+NAA) or an equivalent volume of NaOH (-NAA) as a control. Plates were then scanned using an Epson Perfection V700 scanner.

**Figure 2.**
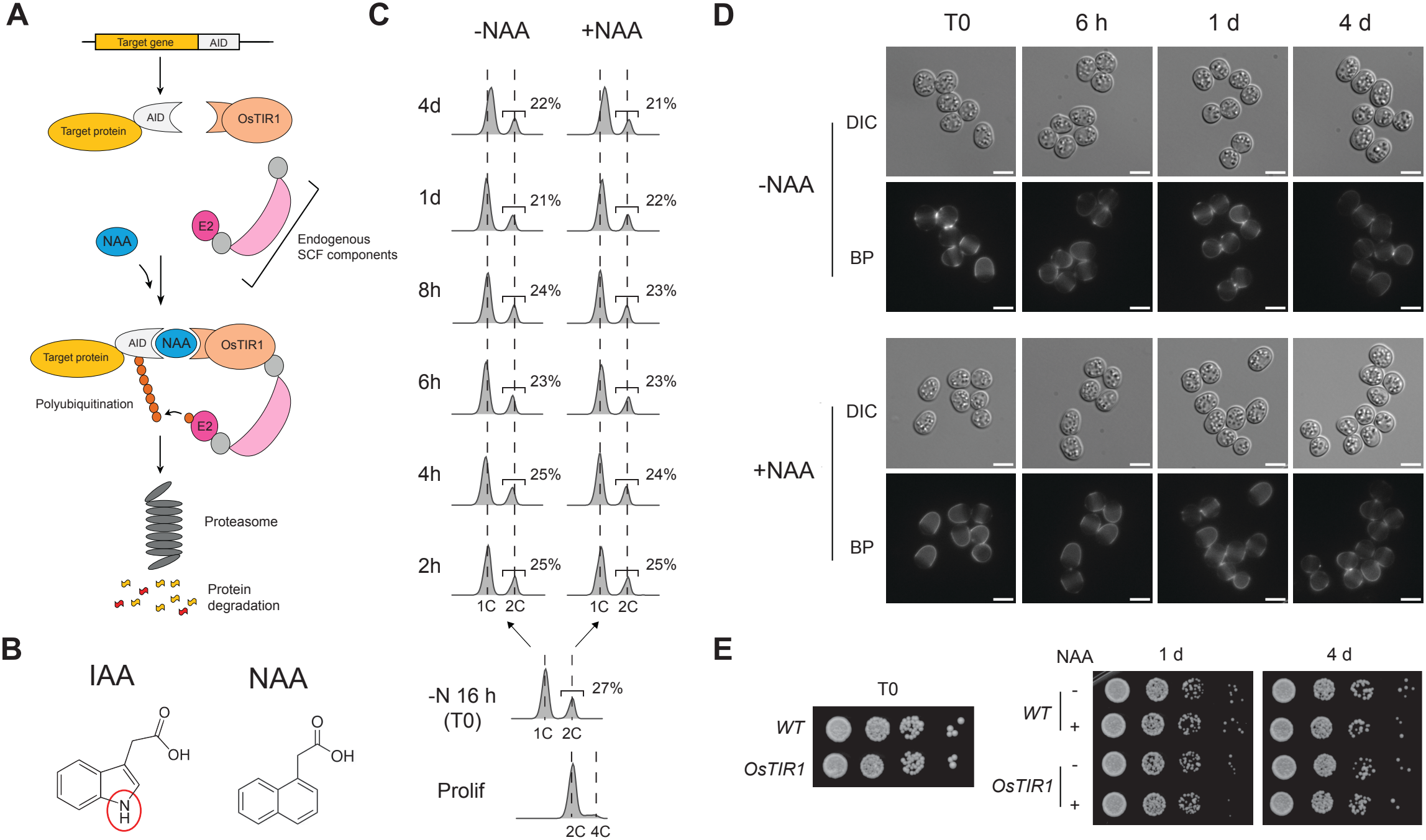
Quiescent cells treated with the synthetic auxin NAA show no cellular defects and do not re-enter the division cycle. **A**. Schematic representation of the auxin-degron system. In the presence of auxin (here the synthetic auxin NAA is shown) and the F-box protein OsTIR1, AID (Auxin-Inducible Degron)-tagged proteins are ubiquitinated and targeted to degradation by the proteasome. SCF: Skp1-Cullin-F-box. **B**. Chemical structure of natural auxin (indole-3-acetic acid, IAA), which contains a nitrogen group (red circle), and synthetic auxin NAA (1-naphthaleneacetic acid), which is devoid of nitrogen groups. **C**. DNA content analysis of nitrogen-depleted quiescent cells expressing the OsTIR1 protein, at the indicated times after treatment with 0.5 mM NAA (+NAA) or with an equivalent volume of the solvent NaOH (-NAA) as a control. Nitrogen starvation and quiescence entry are as described in Fig. 1. Prolif: proliferating cells; -N 16 h: nitrogen-starved quiescent cells. **D**. Representative images of quiescent cells expressing OsTIR1 treated with NaOH (-NAA) or with 0.5 mM NAA (+NAA) for the indicated times. DIC: Differential interference contrast. BP: Blankophor. Scale bar = 5µm. **E**. Quiescent cells of the indicated genotypes were treated with 0.5 mM NAA (+) or an equivalent volume of NaOH (-NAA) and spotted at 0, 1, and 4 days after treatment on YE4S plates to induce cell cycle re-entry.

**Figure 3.**
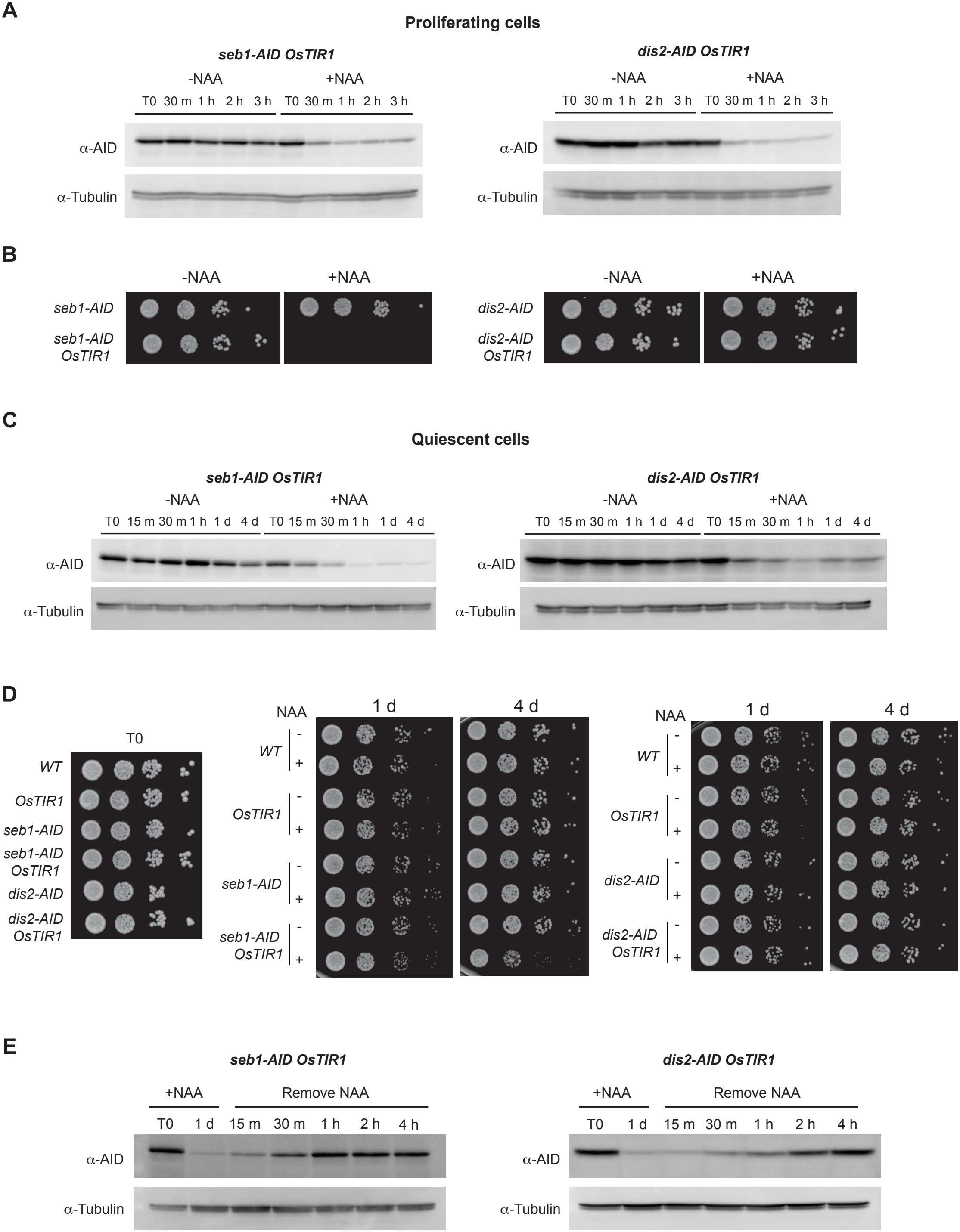
Characterization of the NAA/degron system in proliferating and quiescent cells. **A**. Western blot analysis of protein degradation induced by NAA treatment in proliferating cells. *seb1-AID OsTIR1* and *dis2-AID OsTIR1* cells were treated with 0.5 mM of NAA (+NAA) or an equivalent volume of NaOH (-NAA) for the indicated times. Tubulin was used as a loading control. **B**. Proliferating cells of the indicated genotypes from *A* were spotted on rich medium containing 0.5 mM of NAA (+NAA) or an equivalent volume of NaOH (-NAA). **C**. Quiescent cells of the indicated genotypes were treated with 0.5 mM NAA (+NAA) or an equivalent volume of NaOH (-NAA). Cells were treated at T0 as in Fig. 2C. Protein degradation at the indicated times was probed by Western blot analysis. Tubulin was used as a loading control. **D**. Quiescent cells of the indicated genotypes treated with 0.5 mM NAA (+) or an equivalent volume of NaOH (-NAA) were spotted at 0, 1 and 4 days after treatment on YE4S plates to induce cell cycle re-entry. **E**. Quiescent cells as in *C* were treated with 0.5 mM of NAA (+NAA) or an equivalent volume of NaOH (-NAA) for 24 h. NAA was subsequently washed off, and cells were resuspended in nitrogen-free medium. Protein recovery was assessed at the indicated times by Western blot analysis. Tubulin was used as a loading control.

### Western blot analysis

For each sample, 20 mL of cells at OD = 0.4-0.5 were centrifuged at 3000 rpm for 3 min at room temperature. Cell pellets were washed using 1 mL 20% Trichloroacetic acid (TCA, Sigma-Aldrich), and resuspended in 50 µL 20% TCA prior to freezing at -20°C. Cell breakage was performed using glass beads (Soda Lime Glass beads #11079105 BioSpec products) in a Precellys Evolution (Bertin Instruments). 200 µL of 5% TCA was then added to each sample, and the lysates were transferred into new tubes. Beads were washed with and resuspended in 200 µL of 5% TCA. Lysates were then centrifuged at 3000 rpm for 10 min at room temperature, and pellets were resuspended in 100 µL lysis buffer (300 mM Tris pH 8, 2% SDS), followed by heating for 5 min at 95 °C. Cell lysates were then centrifuged at 14000 rpm for 2 min at room temperature, and supernatants were transferred to a new tube. Protein concentrations were determined using the BCA protein assay kit (ThermoFisher) and diluted in 5X sample buffer (300 mM Tris-HCl pH 6.8, 20% β-mercaptoethanol, 20% SDS, 0.05% bromophenol blue, 25% glycerol) before heating to 5 min at 95 °C. Samples were then loaded in SDS-polyacrylamide gels, and proteins were transferred onto PVDF membranes using the Bio-Rad Transblot Turbo Transfer System. Antibodies were used at the following dilutions: anti-mini-AID-tag (1/1000, MBL M214-3), anti-Tat1 monoclonal (1/20000, a gift from K. Gull), anti-mouse-IgG-HRP (1/100000, Abcam AB97046). Proteins were visualized using an ECL kit (Cytiva RPN2235) using a ChemiDoc MP Imaging System (Bio-Rad).

## Results

### Kinase inhibition using an analog-sensitive approach is unsuitable for nitrogen-starved fission yeast cells

Kinases are critical regulators of a broad range of cellular processes. Analog-sensitive kinase alleles harboring an enlarged ATP-binding pocket allow for specific, dose-dependent and reversible inhibition by bulky non-hydrolyzable ATP analogs (Bishop et al., 2000) (*e*.*g*. 3MB-PP1 (C. Zhang et al., 2013), Fig.1B). This strategy has been applied to a variety of kinases and has proven to be instrumental for the dissection of a number of essential mechanisms (Cipak et al., 2011; Coudreuse & Nurse, 2010; Viladevall et al., 2009). However, the effectiveness of this approach for quiescent fission yeast cells remained unknown.

Surprisingly, we found that treating a population of nitrogen-starved, quiescent wild-type fission yeast cells with 20 µM 3MB-PP1, a concentration commonly used in experimental contexts (Perrot et al., 2018; Viladevall et al., 2009), is sufficient to trigger cell cycle re-entry in a significant fraction of cells. Indeed, we observed that a subset of cells entered S phase within 2-4 h following the addition of the analog (Fig. 1C, red profiles). At 6 h post-treatment, ∼20% of the population had transitioned from 1C to 2C DNA content, suggesting that they underwent genome duplication (Fig. 1C, red profiles). This was followed by cell division (Fig. 1C, blue profiles), with the population returning to the initial proportions of 1C vs. 2C cells within 24 h after 3MB-PP1 addition (Fig. 1C). In addition, we observed significant cell elongation at the 6 h time point, as well as the formation of division septa and clear evidence of cytokinesis (Fig. 1D, 3MB-PP1). These changes in DNA content and cell morphology were not observed in the control population (Fig. 1C-D). Consistent with the DNA content analyses, cell shape and size after 24 h 3MB-PP1 treatment were similar to that at T0, suggesting that a proportion of the population underwent a single division cycle and then exited the cell cycle (Fig. 1D). We note that this cell cycle re-entry remains significantly less efficient than a complete quiescence exit that is brought about by addition of high concentrations of a nitrogen source (Fig. 1A), in which all cells have duplicated their genome after 6 h.

Altogether, our data demonstrate that the addition of micromolar levels of 3MB-PP1 is sufficient for a significant fraction of quiescent cells to resume their cell cycle. We surmise that the minimal amounts of nitrogen present due to the nitrogen groups in the ATP analog (Fig. 1B) trigger growth and cell cycle re-entry. Therefore, in this context, it is extremely complicated to uncouple cellular phenotypes that are associated with the inhibition of a modified kinase from those that result from cell cycle re-entry upon analog treatment. We thus conclude that this strategy is highly inadvisable for this quiescence model, or, at the very least, should be employed with the utmost caution.

### Treatment of nitrogen-starved quiescent cells with the synthetic auxin NAA

As a broader approach for conditional induction of protein depletion in quiescence, we assessed the efficiency of the Auxin-Inducible Degron (AID) system (Nishimura et al., 2009). This method has been successfully used in many organisms, including in the fission yeast, to study protein function in proliferating cells (Kanke et al., 2011; Yesbolatova et al., 2020). The AID strategy relies on tagging a protein of interest with a small peptide sequence that, upon auxin treatment, is recognized by an F-box protein, TIR1 (Fig. 2A). This results in ubiquitination of the target and its subsequent degradation by the proteasome. However, during quiescence, a large spectrum of cellular functions are downregulated (de Morree & Rando, 2023; Marguerat et al., 2012; Sun & Gresham, 2021), and changes in proteasome localization have been observed (Laporte et al., 2008; Takeda et al., 2010). It therefore remained unknown whether AID-mediated protein degradation would be sufficiently effective for functional studies in nitrogen-depleted quiescent fission yeast cells.

Similar to the ATP analog described above, most auxin ligands that have been used to trigger the degradation of AID-tagged proteins, including the natural auxin indole-3-acetic acid (IAA), contain at least one nitrogen group (Fig. 2B). This may also represent a source of nitrogen, thereby making this system problematic for investigating the biology of quiescent cells. Interestingly, the synthetic auxin 1-naphthaleneacetic acid (NAA) is devoid of nitrogen (Fig. 2B) and thus represents a promising candidate for taking advantage of the AID system in nitrogen-starved quiescent cells.

We first tested how quiescent cells expressing an OsTIR1 protein that is codon-optimized for fission yeast, under the control of the *adh1* promoter (X.-R. Zhang et al., 2022), respond to the addition of NAA. We used a final concentration of 0.5 mM, which has been reported to efficiently trigger protein degradation in proliferating cells without affecting cell growth (Kanke et al., 2011; X.-R. Zhang et al., 2022). Flow cytometry analysis of NAA-treated, nitrogen-depleted quiescent cells showed no detectable changes in DNA content during the first hours of treatment and up to four days post-treatment (Fig. 2C). Furthermore, we detected no changes in cell morphology, with quiescent cells maintaining their characteristic small size and rounded shape, and cells did not form division septa (Fig. 2D). We then evaluated the effect of NAA on cell cycle re-entry, taking quiescent wild-type and OsTIR1-expressing cells treated with NAA for 1 and 4 days and spotting them onto rich media. As shown in Fig. 2E, NAA had no effect on the capacity of cells to re-enter a proliferative state. Altogether, these results demonstrate that in contrast to 3MB-PP1, the synthetic auxin NAA has no detectable impact on the physiology of nitrogen-starved quiescent fission yeast cells.

### The NAA/degron system induces rapid, long-term, and reversible protein depletion in quiescent cells

To test the effectiveness of NAA-induced degradation in quiescence, we focused on two proteins: Dis2, a non-essential protein phosphatase, and Seb1, an essential factor that plays a role in poly(A) site selection (Wood et al., 2012). To this end, we tagged both proteins with three copies of a short AID sequence that was shown to improve the efficiency of degradation compared to previously described degrons when combined with the OsTIR1 described above (X.-R. Zhang et al., 2022). In proliferating cells, we observed that both Dis2-AID and Seb1-AID were rapidly degraded within 1 h after NAA addition, with Dis2-AID depletion being slightly more efficient (Fig. 3A). However, *seb1-AID OsTIR1* cells did not grow when spotted on nutrient-rich medium plates containing NAA, showing that the reduction in Seb1 levels is sufficient to impair cell proliferation (Fig. 3B). In contrast, the *dis2-AID OsTIR1* strain did not display growth defects on NAA plates, consistent with Dis2 being non-essential for cell proliferation.

In quiescent cells, Seb1-AID and Dis2-AID were both efficiently degraded within 1 h of NAA addition, with Seb1 dropping to virtually undetectable levels (Fig. 3C). Moreover, both proteins remained at low levels even 4 days after NAA treatment; note that the ligand is not replenished during this time period. These results demonstrate that NAA-induced degradation of AID-tagged proteins in quiescent cells is not only rapid and efficient but also long-lasting. Using a standard spot assay, we then assessed the ability of these cells to re-enter the cell cycle and proliferate in the absence of NAA (Fig. 3D). Loss of Seb1 in quiescent cells resulted in a clear viability defect after 4 days of NAA treatment, with a modest defect being visible after 1 day (Fig. 3C). In contrast, depletion of Dis2 had no effect on growth resumption. Finally, an additional advantage of the auxin-degron system is its reversibility, making it possible to perform transient protein depletion experiments. To evaluate this feature, we treated quiescent cells with NAA for 24 h, then removed the ligand from the culture. This was followed by resuspension of cells in fresh, nitrogen-free medium to maintain their quiescent state. Strikingly, Seb1 and Dis2 levels were restored within 1 h and 4 h after NAA removal, respectively, demonstrating the reversibility of the system even as cells remain in quiescence. Altogether, these results establish the NAA/degron system as a versatile approach for triggering the rapid, long-term, and reversible degradation of proteins of interest in quiescent cells.

## Discussion

In this study, we have probed two different systems for conditional modulation of cellular processes in nitrogen-depleted quiescent cells: 1) a targeted methodology that takes advantage of engineered kinases and non-hydrolyzable ATP analogs for dose-dependent and reversible inhibition of kinase activity and 2) a broadly applicable auxin-degron system for the depletion of AID-tagged proteins of interest.

Our results showed the first approach, while extremely powerful in proliferating cells, is incompatible with nitrogen-starved quiescent cells. Indeed, these cells are exquisitely sensitive to the ATP analog: the addition of low levels of this molecule at concentrations commonly used in experiments was sufficient to promote cell cycle re-entry in a significant fraction of the population. While this does not completely preclude the exploitation of this strategy for assessing the roles of kinases during quiescence, it must be employed with significant caution. In such cases, it may be necessary, on a case-by-case basis, to determine the lowest concentration of ATP analog that is sufficient to inhibit the function of a given altered kinase and subsequently test whether at these levels, the molecule triggers cell growth and division. Nevertheless, it is difficult to eliminate the possibility that even low concentrations of an analog will affect cellular physiology, thus confounding the interpretation of obtained results.

In contrast, treatment of cells with the nitrogen-free synthetic auxin NAA at conventionally used concentrations had no detectable effect on quiescent cells, whether on their DNA content, morphology, or capacity for cell cycle re-entry. Importantly, using two representative targets, an essential and a non-essential protein, we demonstrated that the NAA/degron system mediates efficient, stable, and reversible degradation in nitrogen-starved cells. Moreover, we further showed for the first time that in these non-proliferating cells, not only is this conditionally-induced degradation rapid and reversible, but it is maintained for at least 4 days without ligand renewal. This makes the NAA/degron system an ideal tool for interrogating the mechanisms that govern different stages of the quiescence process, from entry, maintenance, to exit.

## Acknowledgments

We thank members of the Genome Dynamics and Maintenance group for helpful discussions. The fission yeast strain JPNBRP202225 was provided by the National BioResource Project (NBRP Yeast), Japan. This work was supported by grants to PYJW from the Agence Nationale de la Recherche (PRC ExQui, ANR-21-CE12-0025) and the Région Nouvelle Aquitaine (program CHESS, grant agreements 1593520, 1594420, 36167120, and 36155720). We also thank Damien Coudreuse for critical reading of the manuscript.

